# RiboXYZ: A comprehensive database for visualizing and analyzing ribosome structures

**DOI:** 10.1101/2022.08.22.504886

**Authors:** Artem Kushner, Anton Petrov, Khanh Dao Duc

## Abstract

Recent advances in Cryo-EM led to a surge of ribosome structures deposited over the past years, including structures from different species, conformational states, or bound with different ligands. Yet, multiple conflicts of nomenclature make the identification and comparison of structures and ortholog components challenging. We present RiboXYZ (available at https://ribosome.xyz), a database that provides organized access to ribosome structures, with several tools for visualisation and study. The database is up-to-date with the Protein Data Bank (PDB) but provides a standardized nomenclature that allows for searching and comparing ribosomal components (proteins, RNA, ligands) across all the available structures. In addition to structured and simplified access to the data, the application has several specialized visualization tools, including the identification and prediction of ligand binding sites, and 3D superimposition of ribosomal components. Overall, RiboXYZ provides a useful toolkit that complements the PDB database, by implementing the current conventions and providing a set of auxiliary tools that have been developed explicitly for analyzing ribosome structures. This toolkit can be easily accessible by both experts and non-experts in structural biology so that they can search, visualize and compare structures, with various potential applications in molecular biology, evolution, and biochemistry.

## Introduction

The ribosome is a universal RNA-protein complex, whose structure has been a major focus of study for the past decades [1]. Due to the recent progress in imaging technologies and the notable renaissance of cryogenic electron microscopy (cryo-EM) [2], more than a thousand of high-resolution ribosome structures are currently deposited in the protein data bank (PDB) [**?**], encompassing various species, as well as various binding and conformational states [3].

While this variety provides an exciting opportunity to examine specific components of the ribosome through the lens of any single structure available, integrating the data from the PDB for such analysis can be challenging in practice. It is even more challenging to develop computational pipelines that would provide a comparison across multiple structures selected by a user. More precisely, identifying the different components – e.g. rRNA, proteins or ligands – of a given structure can require some manual curation, due to different ontologies of its components. While a ribosome structure-based nomenclature has been proposed [4], the PDB still contains numerous references to earlier naming systems, which lead to unrelated ribosomal proteins being assigned the same name across deposited structures. Ambiguities and inconsistencies are similarly present in the most standard protein databases that include ribosome protein families, such as Pfam [5] or UniProt [6], and have explicitly been mentioned as an obstacle for the ribosome structure-based nomenclature system to be adopted [7].

To facilitate this access and visualization of ribosome components over the whole set of available structures, we present here RiboXYZ (publicly available at https://ribosome.xyz), a database that provides organized access to ribosome structures and enables a direct search and comparison of these structures and their components. In addition, RiboXYZ proposes several export facilities and visualization tools, including the identification and prediction of ligand binding sites, and 3D superimposition of structures and ribosomal components. As illustrated with various examples and applications below, RiboXYZ provides a useful toolkit that complements the PDB database by implementing the current conventions and providing a set of auxiliary tools that have been developed explicitly for ribosomal structures. This toolkit can be easily accessible by both experts and non-experts in structural biology so that they can search, visualize and compare structures, with various potential applications in molecular biology, evolution, and biochemistry.

### Web Server And Database Description

RiboXYZ is a database application that provides organized access to ribosome structures, with several tools for visualisation and export. The user can interact with the database from the application website. The website is mainly divided into two sections, *Database* and *Tools*, whose pages are accessible from the main navigation panel. The database section provides separate access to whole structures, RNA’s, and proteins. The *Tools* section allows to visualize and compare the elements of the database and is composed of three modules which all integrate a 3D viewer [8]: *Visualization, 3D superimposition* and *Ligands/Binding Sites*. All these pages are accessible from the RiboXYZ’s home [page](https://ribosome.xyz/home) after opening a left tab menu. The menu also contains a link to RiboXYZ’s *[How to](https://ribosome.xyz/howto)* page, that contains various tutorials and a user manual providing a complete description of the application and its functionalities, as summarized next.

### General infrastructure and development

RiboXYZ is composed of three services, as shown in Figure 1 **(a)**: the website itself, the Django server and the Neo4j graph-database. The user-facing website is a single-page application (SPA) developed using the React framework for presentational components in front end, and the Redux library for state-management. The Django web-server provides structural computation and exposes the API endpoints. This web-server has access to both the database and static files, with Python libraries (NumPy, SciPy, Biopython) used for most of the computation. The database used is Neo4j and its schema definition language is Cypher, a graph-oriented variant of SQL. Ribosome structures were initially obtained from RCSB PDB‘s Web Service [9], and processed so the structures and their sub-components are nodes of the semantic graph database (as described next in more details). In addition, the database uses [Pfam entries](http://pfam.xfam.org/) [5] to identify ribosomal proteins. The application is deployed to a a *c5a*.*xlarge* instance with 8 GB of memory and 120 GB of disk space storage of the AWS public cloud.

**Figure 1:**
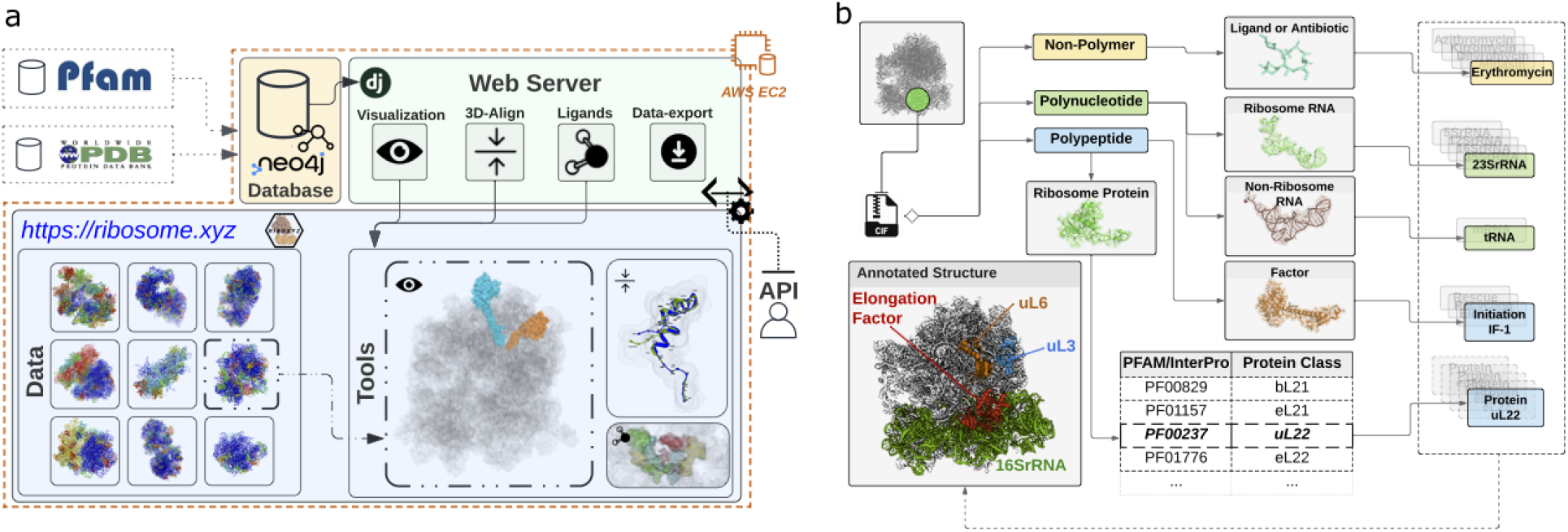
Architecture and data processing of RiboXYZ: **(a)** Diagram that describes RiboXYZ’s architecture. We process data from the PDB and Pfam entries into a graph database structure that is composed of edited structures and their components, which are analyzed for different applications using the web server. The results are mainly accessible to the user via a web interface that give access to both data and visualization tools. **(b)** File processing. Structural files (`.cif`) are processed to first classify chain units. Upon identifying these components, we further edit the chain ID’s of the original structure with our common nomenclature, to facilitate comparison across structures.

### Data structure

Ribosome structures were initially obtained as `.cif` files from the PDB. For each PDB entry, a semantic *profile* of the structure was created and uploaded to the database, that includes annotations provided by the authors. Additionally, external resources associated with the specific PDB entry were obtained as a `.json` file. Individual chains present in each structure are integrated to the database and sorted into various classes, according to the diagram shown in Figure 1 **(b)**. Thus, aside from whole structures, the semantic database includes individual polymer chains, such as proteins and RNA, as well as non-polymer entities (ligands, ions, antibiotics and other small molecules). In particular, more specific attention was given to classifying protein chains, due to the same class of ribosomal protein carrying different names in different structures (for example, ribosomal protein eL21 provided as *60S ribosomal protein L21-A, 60S ribosomal protein L21* or *KLLA0E23651p* in structures PDB 4U4Y, 6LQM and 5IT7 respectively) and, conversely, several instances of ribosomal proteins from different species carrying similar names despite being unrelated in structure and function (like *Ribosomal protein L10 (Predicted)* and *50S ribosomal protein L10* in eukaryotic structure 6R7Q and bacterial structure 5AFI). Although a universal naming system was proposed by a consortium of structural biologists and biochemists [4], applying this consensus retroactively at scale to data already deposited to PDB and elsewhere has seemed prohibitive so far. Thus, our database implements a reduced version of this universal nomenclature upon re-annotating structures from the PDB, by cross-matching a protein’s PFAM families against a manually curated table that maps PFAM clusters [5] to the proposed proteins classes, as illustrated in Figure 1 **(b)**.

### Database content

As of August 2022, the database contains more than 1100 structures that yield ∼ 65000 single protein chains and 5300 RNA chains. Note that the identification of the chains might be missing in most recent structures, due to temporarily incomplete labeling of polymer entities with Pfam entries in the GraphQL endpoint of the RCSB’s data API [9]. Structures were selected from depositions made from 2000, with a resolution that is less than 4 Å (and with the highest resolution obtained so far at 1.5 Å for X-ray structures and 1.98 Å for Cryo-EM). As shown in Figure 2, the number of ribosomal PDB entries has significantly increased over the past decade, with half of the structures of the database having been deposited in the past three years, and more than 90 % from Cryo-EM. In Table, we also detail how these structures are distributed among different species, with more than 59 % of structures accounting for *E. coli, T. thermophilus, S. cerevisiae* and *H. sapiens* as the four major species studied. We anticipate our database to steadily grow and will perform monthly updates to import new depositions from the PDB, via a job scheduler on the server. Also note that as we are preparing the current submission, the Pfam website just announced to be [decommissioned](https://xfam.wordpress.com/2022/08/04/pfam-website-decommission/), which as a result will lead our future updates to rely on Interpro [10], as the source of future Pfam annotations.

**Figure 2:**
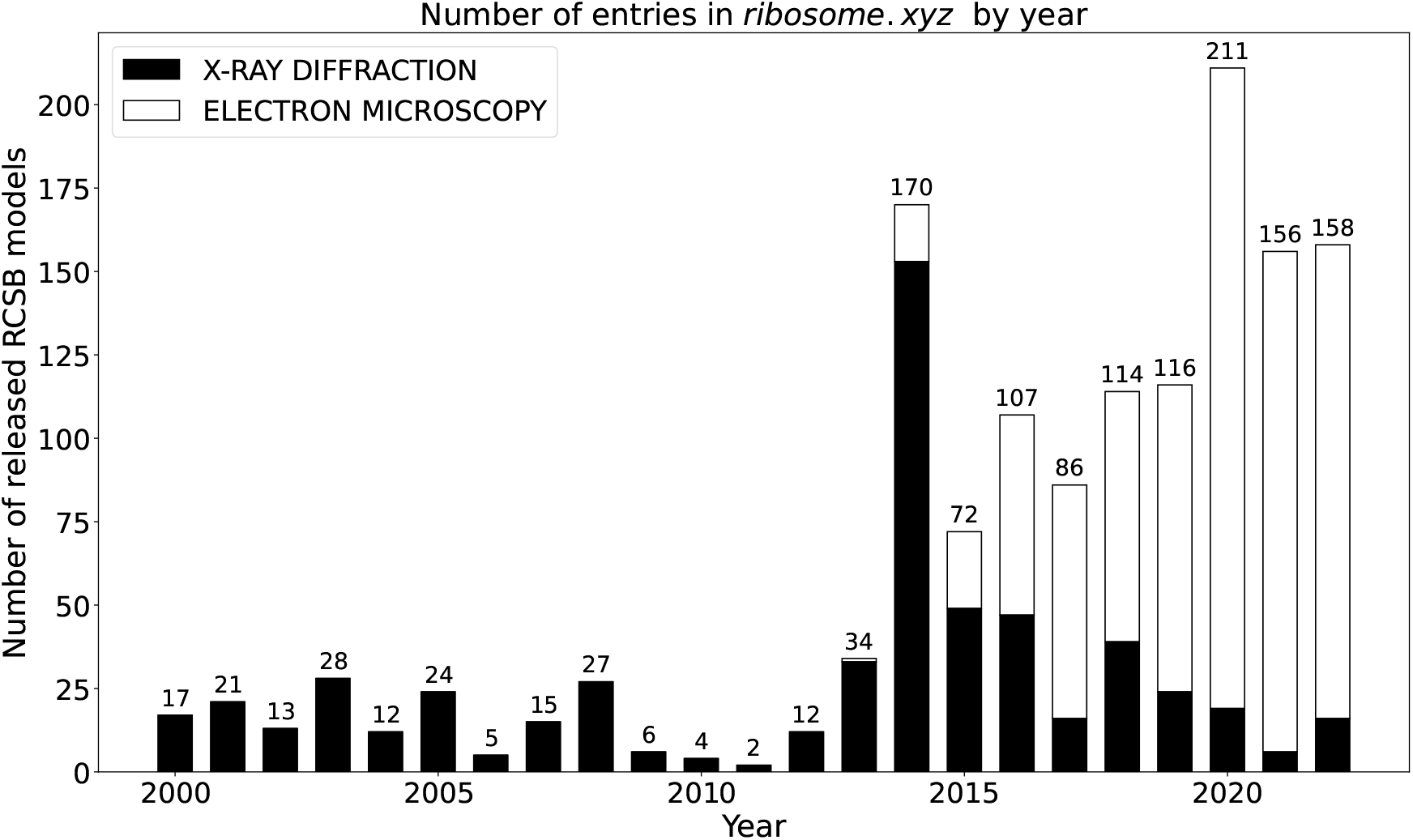
Number of ribosomal structures in the database deposited in the PDB over the past 20 years.

## General features

### Database access and search

The database section is mainly composed of three pages, shown in Figures 3 **(a)-(c)**, that provide separate access to whole structures, RNA’s, and proteins. Each page can be accessed from the website drawer menu, with a set of filters and a search tool bar available to run a search over the database (e.g. by typing species of interest). The pages also contain more detailed information on the structures and their components, and have links to export the data. To facilitate the identification of all the components, the section also contains a specific *Nomenclature* page, shown in Figure 3 **(d)**, that displays the naming system used for ribosome proteins and RNA’s [4]. In addition, this page also provides for each structure the mapping between the protein chain structures ID from the original PDB file and the corresponding protein in the nomenclature.

**Figure 3:**
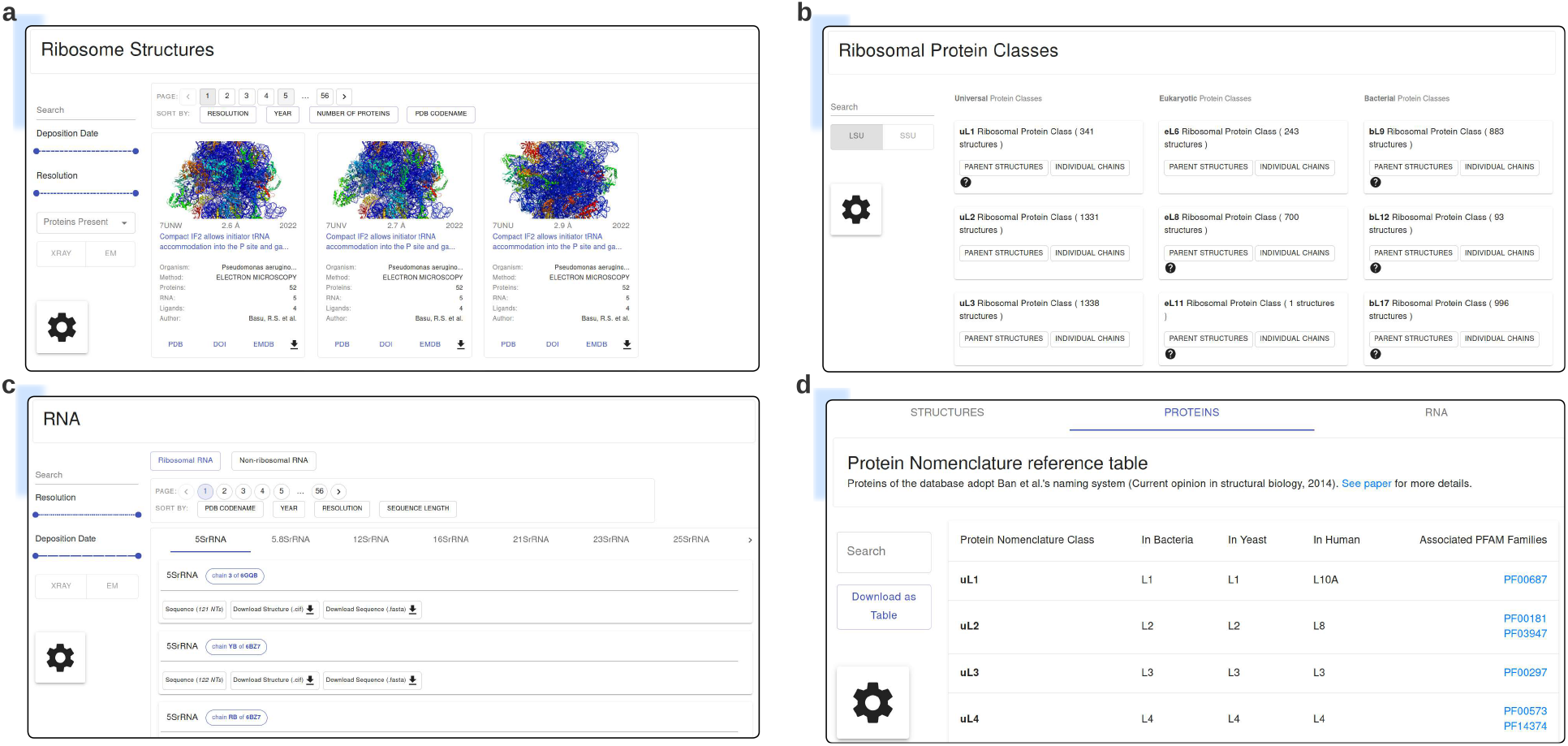
Data section of RiboXYZ, composed of: **(a)** the structure page that provides access to updated ribosome structures (see Figure 1 **(b)**), **(b)** the protein page that covers all the different ribosome proteins, with access to parent structures and individual chains, **(c)** the RNA page that covers all the different RNA components present in the structures of the database (including both ribosomal and non-ribosomal RNA), and **(d)** the nomenclature page that displays the naming system used for ribosome proteins (see also [4] and the *Data structure* section) and RNA’s, and provides for each structure the corresponding mapping with the chain ID’s from the original PDB file.

Each of the structure, protein and RNA pages has specific features. Each structure is represented by an *ID card* containing deposition metadata (e.g. organism, resolution, experimental method, publication info), and links to external resources (PDB, doi, EMDB). Clicking on each ID card displayed on the main screen opens a structure-specific page that lists the structure components. Structures can be downloaded as `.cif` files with chain names edited to match with the standard nomenclature. Each class of ribosomal proteins is also represented by a card, which contains the number of structures in which proteins of this class are present and Pfam annotation. From each class, the user can get access to the parent structures or individual chains. The RNA page has a specific arrangement, separating between ribosomal and non-ribosomal RNA’s, with sub-tabs that correspond with classes (e.g. 5S rRNA, 5.8S rRNA etc. for ribosomal RNA, and mRNA/tRNA for non-ribosomal RNA). Each class links to a list of individual chains associated with the structures present in the database.

### Visualization tools

The *Tools* section allows to visualize and compare the elements of the database. This section contains three separate modules, named *Visualization, 3D superimposition* and *Ligands/Binding Sites*, which can be accessed from the main navigation panel or the database section. We also illustrate each tool in figures 4-6, in the context of some specific examples and applications detailed further below. The tools use the Mol* viewer [8], with multiple features and options to represent the structure (e.g. mapping electrostatic surface potential, solvent accessibility etc.), as well as to annotate and export image. We refer to its [documentation](https://molstar.org/viewer-docs/) for more details. As detailed next, the *Superimposition* and *Ligands/Binding Sites* tools also perform sequence alignment between two given chains, with results that can directly be exported as a `.csv` file.

**Figure 4:**
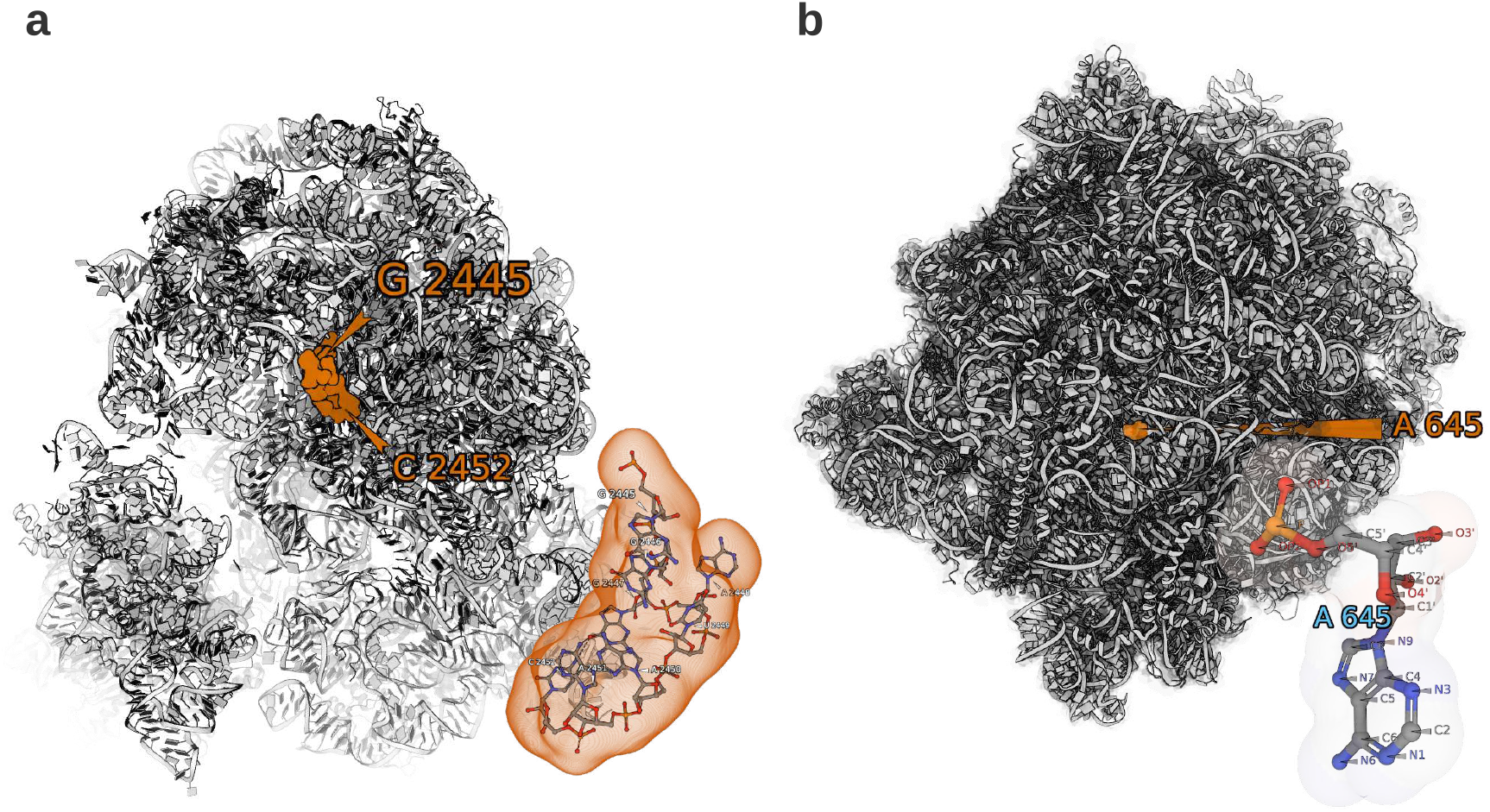
Examples of applications of the *Visualization* tool. **(a)** Localization of the ribosome PTC from PDB 4V7S by selecting nucleotides 2445-2452 from the 23S rRNA. **(b)** Localization of A645 from the 25S rRNA in *S. cerevisiae* (PDB 6QTZ), that is one key single RNA modification site listed in Bailey et al. [14].

Each tool has its own specific features and purpose. The *Visualization* tool allows to select and visualize every single structures, and chain unit (e.g. protein, RNA, ligand) from the database. To do so, a searching tool is also available on the left menu. Once a structure is selected, the user can also highlight any chain, with a further option to select a specific region. Note that while these visualizations can also be done from the RCSB PDB, our database facilitates the search of adequate structure and identification of the sites, by using the same naming nomenclature for the different chain units. The *3D Superimpose* tool executes the [3D superimposition function](https://pymolwiki.org/index.php/Super) `super` from an open source version of [PyMOL](https://github.com/schrodinger/pymol-open-source), to spatially align two chains from two structures of the database. It is similar to the pairwise superimposition tool that was recently added to the PDB, but with a simplified interface and easier access to ribosomal components. The *Ligands/Binding Sites* tool is dedicated to the visualization of components which are neither RNA nor ribosomal protein strands, such as antibiotics, transfer and messenger RNA strands and elongation/initiation factors. These components can be selected and visualized within their parent structures. The tool also allows to predict the potential location of a ligand in any other target structure in which it does not exist. To do so, a binding site is first computed as a list of residues present in the vicinity of the ligand in the original structure (with a cutoff distance set at 5 Å). It is then transposed to the strands of the target structure via Biopython’s `Bio.pairwise2` module [11].

### API

RiboXYZ also allows programmatic access to the data with a dedicated application programming interface (API). The API is partitioned into two endpoints, one for statically-served, downloadable files and the other for querying semantic data from the database. There are multiple endpoints available, with a more detailed description provided in the user manual.

### Examples and Applications

We now provide various examples of applications from RiboXYZ. Some of these examples refer to recent publications and results, which the user can reproduce and interpret by using the database. A more detailed description of the steps associated with each example is provided as tutorials in the database user manual, with videos showing screen recordings. Both the user manual and videos are also directly accessible from the [website](https://ribosome.xyz/howto).

### Residue selection and highlight

To visualize a specific structure or any region of a protein or RNA site of interest, the user can find and select an adequate structure from the database and use the visualization tool, as illustrated in the next two examples. First, we visualize in Figure 4 **(a)** the Peptidyl-Transfer-Center (PTC) region of the ribosome in human, by highlighting the sites 2445-2452 from the *E. coli* 23S rRNA [12]), with all steps to obtain the figure accessible detailed in our online user manual, or by video available from the website. Note that as the PTC is a well-conserved rRNA region [13], the identification of the rRNA site can be done for any species upon aligning its 23S/25S/28S rRNA sequence with that of another well known organism, e.g. human in eukaryote or *E. coli* in bacteria (as further detailed in another example). In our second example (Figure 4 **(b)**), we visualize one of the single rRNA modifications sites (7 sites in 18S rRNA and 73 sites in 25S rRNA) studied in Bailey *et al*., that were recently found susceptible to affect translation, ribosome biogenesis, and pre-mRNA splicing [14]. The figure illustrates the specific location of site 645 of the 25S rRNA, mentioned in Figure 1 of Bailey *et al*..

### 3D superimposition and comparison of homologous proteins

The 3D superimposition tool of RiboXYZ can be used to compare and visualize structural differences of the ribosomal components across different species or conformations or the ribosome (as described in the *Visualization Tools section*). We apply this tool in Figure 5, where we focus on two ribosomal proteins uL4 and uL23. These two proteins are also part of the ribosome exit tunnel, a sub-compartment of the ribosome that contains the nascent polypeptide chain [15]. In particular, comparing and visualizing these two proteins allow to explain and interpret the main differences found in the geometry of the tunnel between eukaryotes and prokaryotes [12], that also impact the nascent chain folding and its escape from the tunnel [16, 17]. In figure 5 **(a)** (with all steps to obtain the figure accessible detailed in our online user manual, or by video available from the website), we illustrate the superimposition obtained for protein uL4 with ribosomes from human PDB 4UG0) and *E. coli* (PDB 5AFI). In particular, we can see an extension of the loop region of the exit tunnel human, that contributes to narrowing down the tunnel at its constriction site, as mentioned in figure 4 of Dao Duc *et al*. [12]. Similar superimpositions can be performed to compare structures across domains of life and species, by finding representative structure in the database (e.g. PDB 4V9F in archaea). In figure 5 **(b)**, we show another comparison between the eukaryotic and prokaryotic ribosomes in uL23, located at the exit port of the tunnel. In bacteria, this region is mainly surrounded by rRNA and the protein uL23, whereas in eukarya and archaea, the segment covering the tunnel region is replaced by the protein eL39 (see also Figure 5 from [12]).

**Figure 5:**
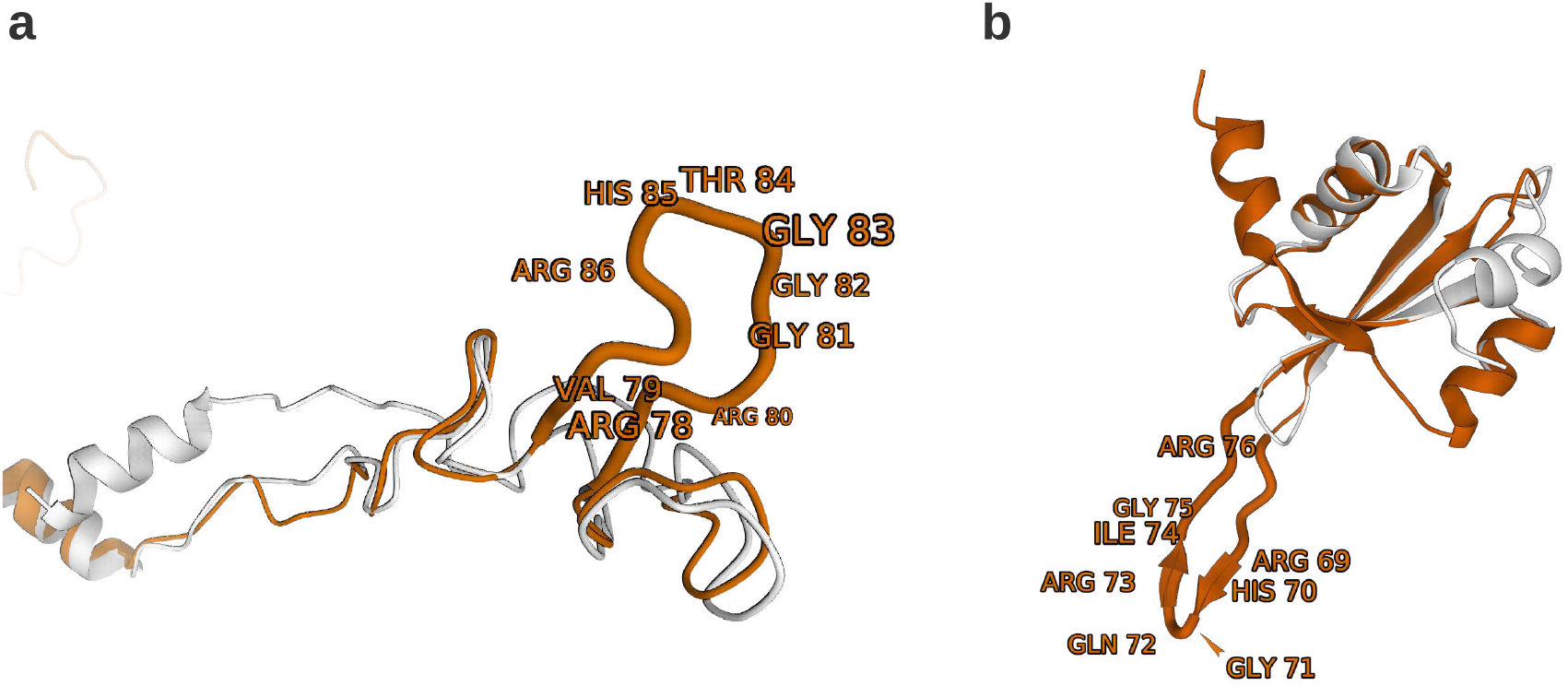
Examples of applications of the *3D superimposition* tool. **(a)** Superimposition of a 90-residue range from proteins uL4 in *H. sapiens* (PDB 4UG0) and *E. coli* (PDB 5AFI), showing the existence of an extended loop at the eukaryotic exit tunnel [12]. **(b)** Superimposition of protein uL23 (same structures as in **(a)**), showing an extra segment in bacteria, replaced by protein eL39 in eukaryote [12]. The eukaryotic proteins are in orange whereas the bacterial are in white and the residues forming the extended loops are highlighted

### Visualization and prediction of antibiotic binding site

The *Ligands/Binding Sites* tool allows to compare and use structures that include ligands (e.g. m/tRNA, translation factors, antibiotics etc.). The user can for example use it to predict and visualize the binding sites of the same ligand across species (assuming that the binding site remains the same), or compare structures from the same organism with and without any given ligand. The prediction of the ligand binding site for an other structure can be useful for molecular docking studies, and is notably relevant for the study of macrolide antibiotics, which constitute an important class of ligands that were found in various structural studies to target different locations of the ribosome, such as the PTC or the exit tunnel [18]. Yet, the structure of a ribosome bound with a specific antibiotic is in general not available for a species that does not belong to the most studied structures (see also Table). For example, the effect of the antibiotic streptomycin on *Mycobacterium tuberculosis* was recently investigated, but with a structure from *Thermus thermophilus* used to visualize the ribosome ligand complex [19], as it is the only available structure of the ribosome bound with streptomycin. We hence first show, in figure 6 **(a)**, how one can use the *Ligand/Binding sites* tool to first visualize the binding site of streptomycin, in the ribosomal structure of the human mitochondrion (PDB 6RW5), and subsequently visualize it in the ribosome of *M. smegmatis* using PDB 5ZEP (Figure 6 **(b)**). To do so, our tool first locates the binding site in 6RW5 at the vicinity of several residues of uS12 (see *Visualization Tools* section), and then produces a list of homologous sites in *M. tuberculosis* from doing pairwise sequence alignment (as described in the *Visualization tools* section), with the result of the alignment shown when clicking on the “Inspect Prediction” button. All the steps to obtain the figure are detailed in our User manual online and accessible by video from the website.

**Figure 6:**
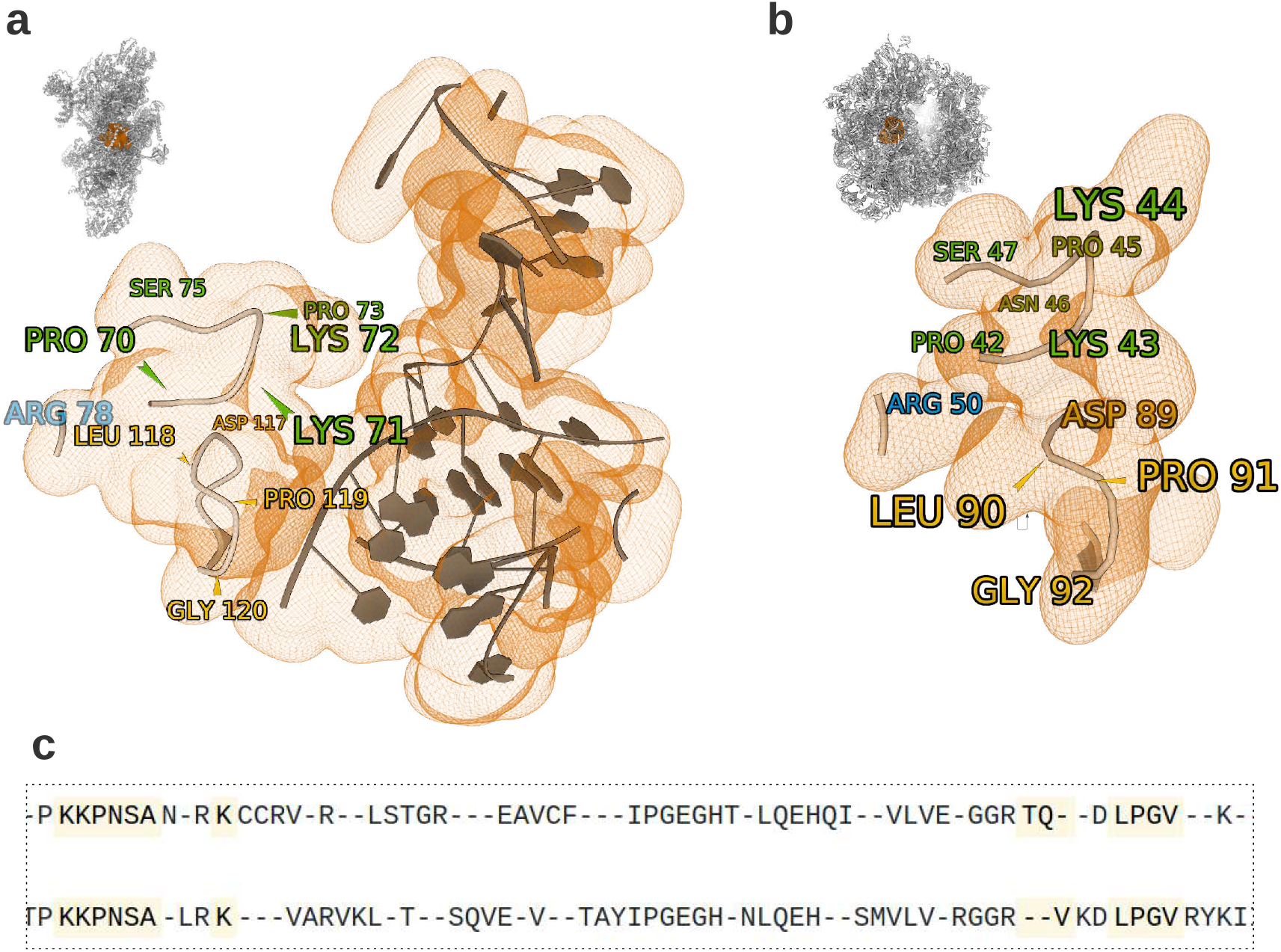
Example of application of the *Ligands/Binding Sites* tool. We use the tool to highlight in **(a)** the structure of PDB 6RW4 (from *T. thermophilus*) with antibiotics streptomycin highlighted, and in **(b)** the prediction of the antibiotic’s binding site in *Mycobacterium tuberculosis*, using the PDB 5ZEP structure (see also [19]). The residues successfully mapped from source structure 6RW4 to target structure 5ZEP are highlighted. **(c)** The sequence alignment between the two uS12 proteins in source and target, on which the prediction is based, as provided by RiboXYZ (using the “Inspect Prediction” button).

### Using curated structures and API

We finally mention two examples of use of the database and its edited structures. First, the database can facilitate the localization of the PTC and exit tunnel for a given set of species. Upon downloading any structure of interest from the database, the user can take advantage of the common ontology used for the RNA and ribosomal proteins, and programmatically select the nucleotides located at the PTC. The tutorial section of our user manual contains a step-by-step description of how to run this procedure. In particular, this procedure can potentially automate the manual processing done in our previous paper [12], to initialize an algorithm extracting the exit tunnel location [20]. While this comparison was done for twenty structures in [12], using RiboXYZ would allow in principle to extend the localization of the tunnel to all the structures that contain the exit tunnel.

Secondly, the webserver ProteoVision [21] provides an example of how RiboXYZ’s API can be used for any application that requires a large number of curated structures. The DESIRE mode of ProteoVision dedicated to evolutionary analysis and visualization of ribosomal proteins (rProteins) across the Tree of Life, provides an integration of the multiple sequence alignments of rProteins with their topology diagrams, and the selected ribsoomal 3D structure. The integration is facilitated by use of API annotations of rProteins from RiboXYZ that resolves the relationship between the protein name according to the current notation [4] and provides a list of all available PDB structures containing the selected protein name. The details of integration are available in the ProteoVision’s github [repository](https://github.com/LDWLab/DESIRE/).

## Discussion

Here we present RiboXYZ, a database that provides a simplified and curated access to ribosome structures with several visualisation tools. The database provides a unified nomenclature of all ribosomal components across all structures available in PDB. While the RCSB PDB server has recently integrated more tools for visualization and comparison [22], the interface of RiboXYZ was designed to be light and intuitive enough to allow non-specialists in structural biology who are unfamiliar with the more dense PDB interface, to easily get access to structures and visualize any result that can relate with some aspect of the ribosome structure, as illustrated in our examples and applications [12, 14, 19]. In this regard, a fundamental contribution of our database is that our data structure disambiguates the identification of the ribosomal components, that was so far hindered by the use of arbitrary ID’s in deposited structures and other naming convention outside from the field of structural biology [5, 6]. Previous efforts to integrate ribosome structural data were made to build databases and interfaces for 3D alignment [23], or jointly visualize 1D, 2D and 3D structures of the ribosome [24]. While these databases are not available anymore and/or do not scale up to the current amount of available structures [3], the past year has interestingly seen the release of new applications with dedicated online server and database, that perform some advanced analysis of ribosome structures and proteins using a large dataset of structures available [21, 25]. For instance, Radtool [25] evaluates the relative rotation of the two ribosomal subunits across all structures available from the PDB. Proteovision, which was earlier mentioned as it uses RiboXYZ’s API allows to perform an evolutionary analysis and visualization of ribosomal proteins over a large number of species in the tree of life. Interestingly, ProteoVision jointly integrates a 3D viewer [8] using structures from the PDB like RiboXYZ, which demonstrates how structure visualization can help interpret results derived from sequence analysis.

The database is also built to be augmented and maintained over time, with the release of new structures, and the potential integration or crosslinking with other databases to improve its coverage and annotations [10]. As RiboXYZ applies a generic framework to process and describe all the structures, some more effort can be done in the future to provide more specific annotations and tools to cover some fundamental aspects of the ribosome that were recently adressed in structural studies, including the conformational heterogeneity [3], assembly pathways [26], and other functions of the ribosome (e.g. antibiotic resistance [18], nascent chain interactions [15]). While the tools produce satisfactory results in our examples, we also plan for further improvement and alternative options. For instance, an alternative to the sequence alignment performed in the superimposition and binding site tools can be brought using pairwise structural alignment methods [27], or by leveraging curated seed-sequences for multiple sequence alignment [21]. Overall, these potential modifications can be easily implemented, given the modular architecture of the database. More generally, while the current tools mostly apply to single structure for visualization, or pairs of structures for superimposition and binding sites prediction, it would be interesting to fully leverage the diversity of structures in our database by designing tools that perform multiple and high throughput analysis across the database.

## Acknowledgements

This research was supported by a NSERC Discovery Grant (PG 22R3468), a UBC STAIR grant, the Peter Wall Institute for Advanced Studies and Biotalent Canada to KDD and AK. This research was also partially supported by the National Aeronautics and Space Administration grant 80NSSC18K1139 to ASP. We thank Loren Williams’s group for discussion.

## TABLES

**Table 1:** Structures in the database sorted by species/domain of life.

| Species | Domain | Total<br>Count | X-Ray | EM | Average resolution<br>(X-Ray/EM) in Å |
| --- | --- | --- | --- | --- | --- |
| *E. coli* (K-12) | Bacteria | 265 | 45 | 220 | 3.2 / 3.3 |
| *T. thermophilus* | Bacteria | 230 | 221 | 9 | 3.1 / 3.6 |
| *H. sapiens* | Eukarya | 103 | 0 | 103 | NA / 3.2 |
| *S. cerevisiae* | Eukarya | 81 | 33 | 48 | 3.1 / 3.5 |
| *H. marismortui* | Archaea | 65 | 65 | 0 | 2.8 / NA |
| Others | All | 393 | 76 | 317 | 3.2/3.2 |

